# ICG-Functionalized Gold Nanostars As An Effective Contrast Agent For Real-time Tumor Localization with Dynamic Optical Contrast Imaging (DOCI) and Enhanced Radiation Therapy

**DOI:** 10.1101/2023.09.19.558473

**Authors:** Yazeed Alhiyari, Yang Liu, Laith Mukdad, Ren A. Odion, Lauran K. Evans, Ramesh Shori, Tuan Vo-Dinh, Maie St. John

## Abstract

The primary management of head and neck squamous cell carcinoma relies on complete surgical resection of the tumor. However, the establishment of negative margin complete resection is often difficult given the devastating side effects of aggressive surgery and the anatomic proximity to vital structures. Positive margin status is associated with significantly decreased survival. Currently, surgeons determine where the tumor cuts are made, by palpating the edges of the tumor and using prior imaging. After a tumor is presumed to be removed in its entirety, the surrounding tissues are sampled by frozen section histologic pathology to ensure that no microscopic disease is left behind, the efficacy of which varies and is subject to sampling error. The methodology by which frozen sections are collected whether tumor bed driven or specimen driven can also alter margin outcome. Thus, improving intraoperative detection of tumor margins is key to optimizing treatment and outcomes. Our group has developed a possible solution for this unmet clinical need. We have previously designed Dynamic Optical Contrast Imaging (DOCI), a novel imaging modality that acquires temporally dependent measurements of tissue autofluorescence. Furthermore, we demonstrated that DOCI can distinguish HNSCC from adjacent healthy tissue with a high degree of accuracy. DOCI images are captured in real time and offer an operatively wide field of view. With the addition of ICG conjugated gold nanostars (GNS) we can improve DOCI image contrast between tumors vs normal tissues as well as use the GNS for CT imaging and radiotherapeutic treatment.

## Introduction

Head and neck squamous cell carcinoma (HNSCC) is the sixth most common cancer in the world. ^1^ The primary management of HNSCC relies on complete surgical resection of the tumor. However, establishing negative margin complete resection is often difficult given the devastating side effects of aggressive surgery and the anatomic proximity to vital structures. Positive margin status is associated with significantly decreased survival, and negative margins are vital to a complete oncologic resection. ^2,3^ Currently, the surgeon’s fingers determine where the tumor cuts are made, utilizing pre-operative imaging and palpating the tumor edges. After a tumor is presumed to be removed in its entirety, the surrounding tissues are often sampled by frozen section (FS) histologic pathology to ensure that no microscopic disease is left behind, the efficacy of which varies and can be subject to sampling error. Multiple studies have demonstrated that FS utilization does not improve survival or locoregional control and, furthermore, is not predictive of final margin status.^3-5^ The methodology by which frozen section margins are collected—whether from the tumor bed or specimen itself—can also alter margin outcome.^6,7^ Thus, improving intraoperative detection of tumor margins is critical to optimizing oncologic surgical treatment and outcomes.

Our group has developed a possible solution for this unmet clinical need. We have previously designed Dynamic Optical Contrast Imaging (DOCI), a novel imaging modality that acquires temporally dependent measurements of tissue autofluorescence.^5-9^ Furthermore, we demonstrated that DOCI can distinguish HNSCC from adjacent healthy tissue with a high degree of accuracy.^5^ DOCI images are captured in real time, offering an operatively wide field of view. Thus, the DOCI system can effectively discriminate cancer from other surrounding tissues in patients and could be utilized in driving additional biopsies or re-resection in real-time when patients are undergoing surgery.

After surgical resection is complete, many patients with HNSCC will undergo adjuvant radiotherapy (RT) with or without chemotherapy or immunotherapy, depending on the characteristics of the tumor. RT to the head and neck region carries a significant decrease in quality of life, causing dysphagia, mucositis, trismus, xerostomia, and tooth decay, with higher dosages increasing the risk of devastating complications such as osteoradionecrosis; thus, decreasing RT dosage to normal tissue while still effectively eradicating malignancy, is a desirable outcome and currently under study.^10^ Dose de-escalation in HNSCC is explored in the current study, via leveraging the mechanics of nanomedicine.

Nanomedicine provides an attractive means for improving DOCI tumor identification, particularly when determining whether additional biopsies or re-resections are needed.^11,12^ Our group has developed a novel surfactant-free method to synthesize biocompatible star-shaped gold nanoparticles, gold nanostars (GNSs).^13^ The synthesized GNSs have tip-enhanced plasmonics in the near-infrared region and are toxic-surfactant free, which is suitable for in vivo biomedical applications. GNSs have been designed to have surface plasmon resonance in the near-infrared (NIR) region, which is the optimal optical window for DOCI due to the low optical attenuation coefficients of water and other tissue chromophores in the 700-1100 nm spectral range.^11,14-16^ We have demonstrated that our GNS nanoparticles can selectively accumulate in tumors but not surrounding healthy tissues due to the enhanced permeability and retention (EPR) effect using brain tumor, bladder cancer, and sarcoma murine animal models.^11,17,18^ GNSs have been used as contrast agents and can be functionalized with a wide variety of molecules suitable for surface-enhanced Raman spectroscopy (SERS), positron emission tomography (PET) imaging, and two-photon photoluminescence (TPL) imaging.^11-13,17,19^ Moreover, prior studies have not only indicated high tumor uptake of GNSs, as well as deep penetration into tumor interstitial space, but also demonstrated effective tumor margin delineation with the use of optical imaging.^11,12,17^ We have additionally performed long-term toxicity study and experiment results demonstrated that our GNSs nanoparticles are biocompatible and have no deleterious effects 6 months after systemic administration.^17^

In the current study, we demonstrate the combined use of GNS technology with DOCI imaging in a tissue model and an in vivo head and neck cancer mouse model, to identify malignant tissue in a deeper tissue plane than DOCI alone, and guide tumor resection margins. This proof-of-concept study demonstrates that GNS technology in combination with DOCI imaging has the potential to revolutionize cancer care by allowing the surgeon to precisely determine cancer margins intraoperatively. The addition of GNS technology in refining DOCI cancer margin delineation can then serve as a platform for multi-center trials for the detection of tumor margins in the head and neck and throughout the body. We additionally illustrate GNS’s potential in RT dose de-escalation in our in vivo model and demonstrate utilization of this compound as a contrast agent. ICG-GNS can thus be employed in multifaceted approaches to cancer treatment: pre-operative visualization, intra-operative real-time margin assessment, and post-operative RT.

## METHODS

### Dye-conjugated Gold Nanoparticle Synthesis and Characterization

Star-shaped gold nanoparticles (GNSs) were synthesized using our developed surfactant-free method.^12^ Synthesized GNSs solution was condensed and purified with 4000 g centrifugation for 1 hour. The pellet was resuspended in 10 mM sodium phosphate buffer at pH 6. SH-PEG-NHS (MW 5000) linker molecules were added to the resuspended GNSs solution with a GNS-to-Linker ratio of 1:10,000. The mixed solution was incubated at room temperature for 1 hour and then purified with 4000 g centrifugation for 1 hour. Indocyanine green (ICG)-Amine molecules were added to the linker-functionalized GNSs solution with an ICG-Amine-to-GNSs ratio of 1:10,000. 50 mM sodium borate buffer (pH 8.5) was added to the reaction solution and incubated for 8 hours. The reaction solution was condensed with centrifugation at 4000 g and then purified by using dialysis. Gold mass concentration for the GNSs solution was measured with inductively coupled plasma mass spectrometry (ICP-MS).

### Tumor Phantom Fabrication

To evaluate the spatial extent of the DOCI+GNS system’s capabilities, we created a series of ICG-GNS 3 w/v% agarose (Sigma Aldrich, New Jersey, US) gel phantoms containing ICG-GNS, as well as liquid ICG-GNS serial diluted in 30ul wells. Such phantoms were formed from a 3D printed rectangular mold of 1.5 cm x 1.5 cm size.

### Dynamic optical contrast imaging instrumentation

The instrumentation and setup for the DOCI system have been previously published.^15^ The DOCI system utilizes a 365-nm UV LED as an excitation source to produce contrast between fluorophores of different decay rates. Fundamentally, DOCI relies on the fact that the longer lifetime fluorophores produce more signal than the shorter lifetime fluorophores when referenced to their steady state fluorescence. To further assist with differentiating the subtle autofluorescence signal indicative of different tissue types, the autofluorescence signal is captured using different spectral filters (bands/channels). To visualize ICG-GNS localization in tissues, a 785nm light source was added to the existing DOCI setup. Both 365-nm and 785nm light source pulses were aligned with both 365nm and 785nm imaging taking place concurrently. Band pass filters were used to spectrally select the emitted light.

### DOCI image processing and statistical analysis

DOCI image processing and calculation has been detailed previously by our group.^5^ In brief, a DOCI image was calculated for 9 filter channels spanning the 400nm-700nm range, and an intensity image is computed for 9 BPF channels. With the addition of 785nm light source for ICG-GNS excitation band pass filters were extended to 840nm. An aggregate intensity measurement was recorded from the moment the illumination started to decrease until the fluorescence ceased. A DOCI image was calculated by normalizing the aggregate fluorescence decay intensity by the aggregate stead-state fluorescence intensity. Data analysis was performed using MATLAB R2021b and Microsoft Office Excel. For each tissue sample, a region of interest (ROI) corresponding to a tissue type identified on frozen histology was selected. The average DOCI value for each ROI was subsequently calculated. DOCI values for each tissue type were normalized against the tumor DOCI value. After confirming a normal distribution, a one-sample t-test was utilized for statistical analysis between different tissue types and cancer.

### In vivo murine model

The use of mice for this study was approved by the Institutional Animal Care and Use Committee (IACUC) at the University of California, Los Angeles. All methods were carried out in accordance with relevant guidelines and regulations, and all methods are reported in accordance with ARRIVE (Animal Research: Reporting In Vivo Experiments) guidelines. Sixteen C3H/HeJ male mice underwent subcutaneous, bilateral flank injection with 500,000 SCC7 cells (RRID:CVCL_V412), a murine head and neck cancer cell line. Tumors were allowed to grow for approximately 2-3 weeks until tumor volume reached 1 cm^3^. 40 tumors were successfully grafted to the hosts and split between all experimental arms (Dose de-escalation, PET/CT imaging and DOCI imaging). Eight mice received 100 μL of ICG-functionalized GNS (80 mg/kg gold) in 1X phosphate buffered saline (PBS) solution injections intravenously via the tail vein using a 30-gauge needle. Two mice received both tail vein ICG-GNS and intratumoral injections. Six mice received intertumoral ICG-GNS injections exclusively. Four mice received no ICG-GNS injections. 24 hours after injection, the mice were anesthetized with weight-based dosing of ketamine/xylazine. The mice first underwent DOCI imaging prior to incision in order to assess depth of imaging through skin. Afterward, under anesthesia, the mice were prepped in a sterile fashion for surgery and a midline incision was made through the skin; bilateral skin flaps were raised until the tumor was exposed. At this point, the mice underwent DOCI imaging to determine the extent of resection necessary. After assessing DOCI imaging, PET-CT imaging was done at 12 and 48 hours post tail vein injection. Both tumor and adjacent tissue margins were sent to the UCLA translational pathology core laboratory (TPCL) for permanent sectioning. In subsequent dose de-escalation experiments eight mice received tail vein injection of ICG-GNS 150 μL of ICG-functionalized GNS (80 mg/kg gold) which were split into two groups of 4 for radiotherapy, one group received 4x2Gy and the other 4x4Gy respectively. Four mice had a total 100ul of (80 mg/kg gold) ICG-GNS intratumorally injected evenly through the 4 quadrants of the tumors and received 4x4Gy of RT. Four mice were given intratumoral ICG-GNS without RT, and the last four mice received 4x4Gy RT with no administration of ICG-GNS.

### CT and PET-CT Imaging

To further investigate GNSs dynamic biodistribution and to display intertumoral accumulation, we performed high resolution CT and dual-energy CT imaging on all sixteen mice with primary SCC7 tumors 48 hours after ICG-GNS injections. Mice with primary head and neck SCC are immunocompetent and may more accurately reflect the tumor microenvironment and response to therapy of human tumors compared to xenograft models.^20^

### Radiation therapy and tumor growth dynamics

All sixteen mice with tumors were anesthetized with ketamine/xylazine and had eye ointment applied to their eyes prior to radiation therapy (RT). The mice were positioned under ½ inch of lead shielding leaving on the tumors exposed. An X-ray dose was delivered at 0.4299 Gy/min for 9.3 min until 4 Gy of total dose was received. Mice received either a standard 4 x 4 Gy RT, which is the scaled comparable dose given to head and neck cancer patients, or a de-escalated dose of 4 x 2 Gy RT.^21,22^ To assess overall changes in tumor volume following radiation therapy treatment, each tumor was measured across their length, width, and height using calipers. Tumor volume was measured daily over a course of 3 weeks and calculated as *Tumor volume* = *π/6* × *length* × *width* × *height*.

## RESULTS

### DOCI and Fluorescence Intensity Measurements of Phantom Models

We evaluated the sensitivity and established the limit of detection in tissue phantom models using ICG-GNS. To evaluate the spatial extent of the DOCI+GNS system’s capabilities, we plated 15ul of ICG-GNS and Rhodamine B (RhB) in 30ul wells for DOCI imaging. Fluorescence was measured across this matrix of parameters to form a basic calibration for depth sensitive detection from the set of vertical offsets from the surface excitation and collection spot. In **Figure 1**, we noted that that with dual simulations excitation (365nm/785nm) RhB illuminates at the 540nm BPF (30nm wide) and 800nm LPF (long pass filter) while ICG-GNS illuminate at the 800nm LPF exclusively. This is consistent with the emission profiles established through literature and plotted on the right side of figure 1. Evaluating the DOCI machines sensitivity to the concentration injected in mice we found that ICG-GNS concentrations of 2.5mg/ml and 0.25mg/ml give the same DOCI value of 0.4. Lower concentration of ICG-GNS results in a rise in the DOCI value to 0.43 while PBS vehicle control had a DOCI value of 0.48 (p < 0.5).

**Figure 1:**
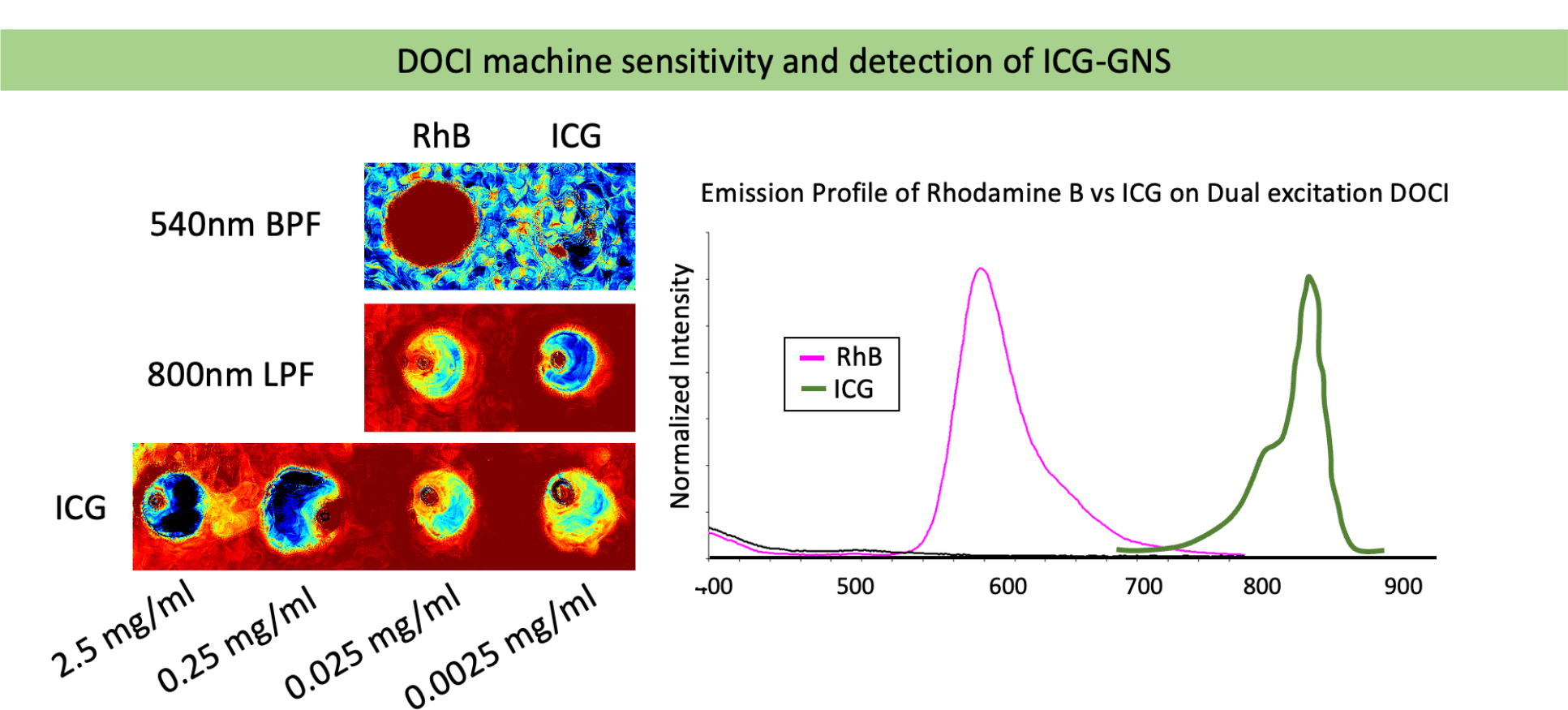
Left illustrates Rhodamine B (RhB) vs ICG after dual excitation (365nm/785nm) filter gating at 540nm BPF. RhB is visible but ICG is not. At 800LPF both RhB and ICG are visible. Lastly, DOCI lifetime measurement changes at concentrations below 0.25mg/ml. The right panel depicts emission spectra for RhB and ICG.

### DOCI Measurements of Murine Models

After calibration with phantoms, an orthotopic H&N cancer murine model was utilized to localize the spatial positions of the ICG-GNS in the tumor itself. The ICG-GNS was tuned to have maximal contrast enhancement for DOCI at 800-840 nm with 785nm excitation light source, and thus would not interfere with traditional 365nm DOCI signal occurring in the 400-790nm range. **Figure 2** shows 3 mice: The first mouse has not been exposed to any ICG-GNS treatment; the second mouse underwent intratumoral injection in each of the four quadrants of the tumor, receiving a total dose of 100ul of 80mg/ml ICG-GNS; and the last mouse received 100ul of 80mg/ml ICG-GNS tail vein injection. Tumors were excited with both 365nm/785 dual-excitation via the DOCI machine. Intratumorally and tail-vein injected tumors demonstrate ICG-GNS patterns unique to their treatment administration type under the 800nm LPF. Tail-vein injected mice accumulated ICG-GNS around the tumor margins, while intratumorally injected tumors concentrate ICG-GNS at injection site locations. Notably, ICG-GNS does not improve DOCI imaging unless excited by a 785nm light source. **Figure 3** demonstrates DOCI imaging with both 365nm and 785nm light sources. The 785nm beam penetrates deeper into the tissue to excite ICG-GNS. **Figure 3 panel 3** demonstrations that UV cannot identify tumor tissue beneath the skin; however, a tumor that has received ICG-GNS can be adequately visualized with 785nm excitation and 800nm LPF (**Figure 3 panel 4**).

**Figure 2:**
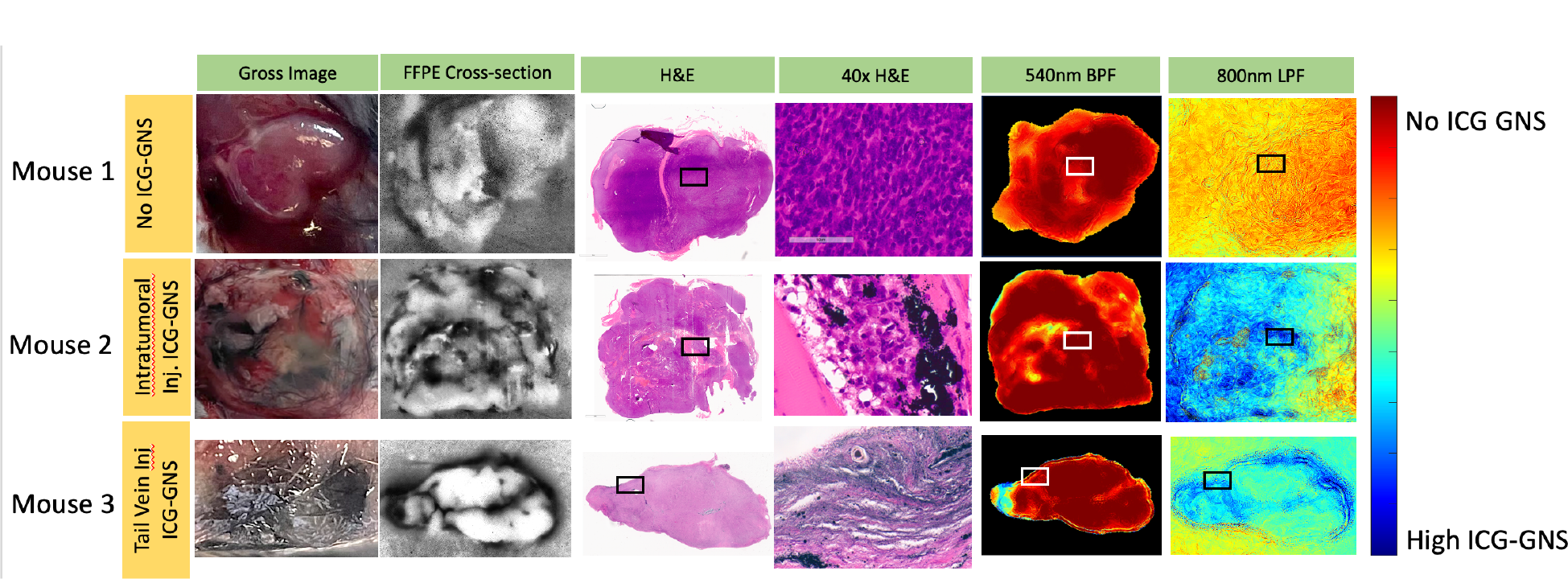
Depicts a mouse that received no ICG-GNS (Mouse 1), intratumorally injected ICG-GNS (Mouse 2) and tail vein ICG-GNS injection (Mouse 3). Each black or white box depicts location of 40x zoom on H&E panel. Tumors were resected from mice, cut in cross-section, formalin fixed, and paraffin embedded for H&E staining. Tumor were imaged with dual excitation (365nm/785nm) and filter gated at 540nm and 800nm. Results shows the “blue” coloring on that 800nm LPF correlate with location of ICG-GNS accumulation as depicted on 40x H&E slides.

**Figure 3:**
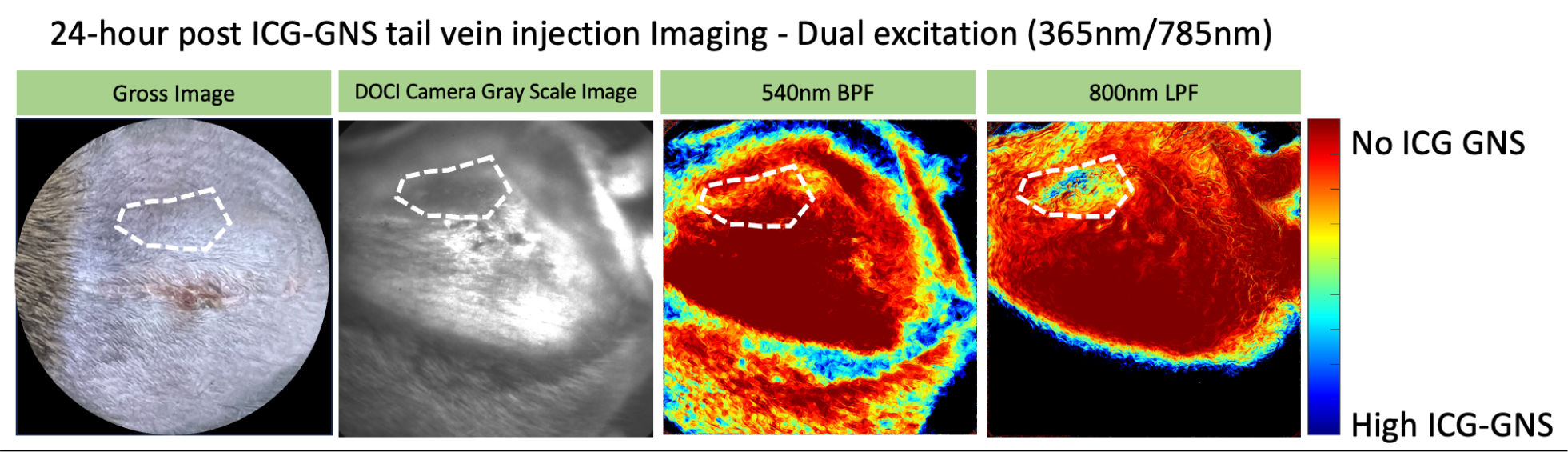
Depicts selective ICG-GNS detection under the skin of a mouse with a tumor deep to the skin, as demonstrated in the 800nm LPF, but not the traditional 540 BPF. This mouse had undergone ICG-GNS tail vein injection 24 hours prior to the above DOCI imaging.

### Histological Evaluation of ICG-GNS Injected Tumors

To evaluate the localization and distribution of ICG-GNS, tumor specimens were collected post imaging and processed for hemotoxin and eosin (H&E) staining. Although ICG-GNS is not characterized to be absorbed or permanently intercalate into the tissues, ICG stain could be observed in both intratumorally injected and tail-vein injected mice. **Figure 2, columns 3 and 4** demonstrates that intratumorally injected ICG-GNS was found in very clumped patches throughout the tumor or localized around nucleated cells in smaller concentrations. **Figure 2 columns 4** shows at 40x magnification several patches of ICG-GNS. Tail vein injected mice showed ICG-GNS accumulation along the tumor margins, however, at much lower concentrations then intratumorally injected mice. After tumors were removed, residual tumor beds were evaluated for the presence of ICG-GNS, and no identifiable ICG-GNS could be localized via histology.

### CT and PET-CT Imaging

As previously published in our prior studies,^11-13^ tail vein injection of ICG-GNS results in higher uptake and imaging contrast within the tumor and liver within our murine model (**Figure 4**). PET-CT imaging expectedly indicated FDG activity within the region of the flank tumors. Intertumoral ICG-GNs injection demonstrates sharp increase contrast at injection sites on CT imaging. The CT value increases from 80HU for tumor without ICG-GNS, to 89HU for tumor with intratumoral injection, and 87HU for tumor with tail vein injection. Using volumetric region of interest ROI on the total tumor, tumor HU values were considered significant over non-inject tumors. This could be accounted by the fact that tail vein injected tumors only show increased contrast on the tumor edge and not thought out the whole tumor, while intratumorally injected tumors only demonstrated areas of high contrast at injection sites, and not throughout the tumor as well. Thus, tumors with both intertumoral and tail-vein injections may significantly average in pixels with little or no GNS uptake.

**Figure 4:**
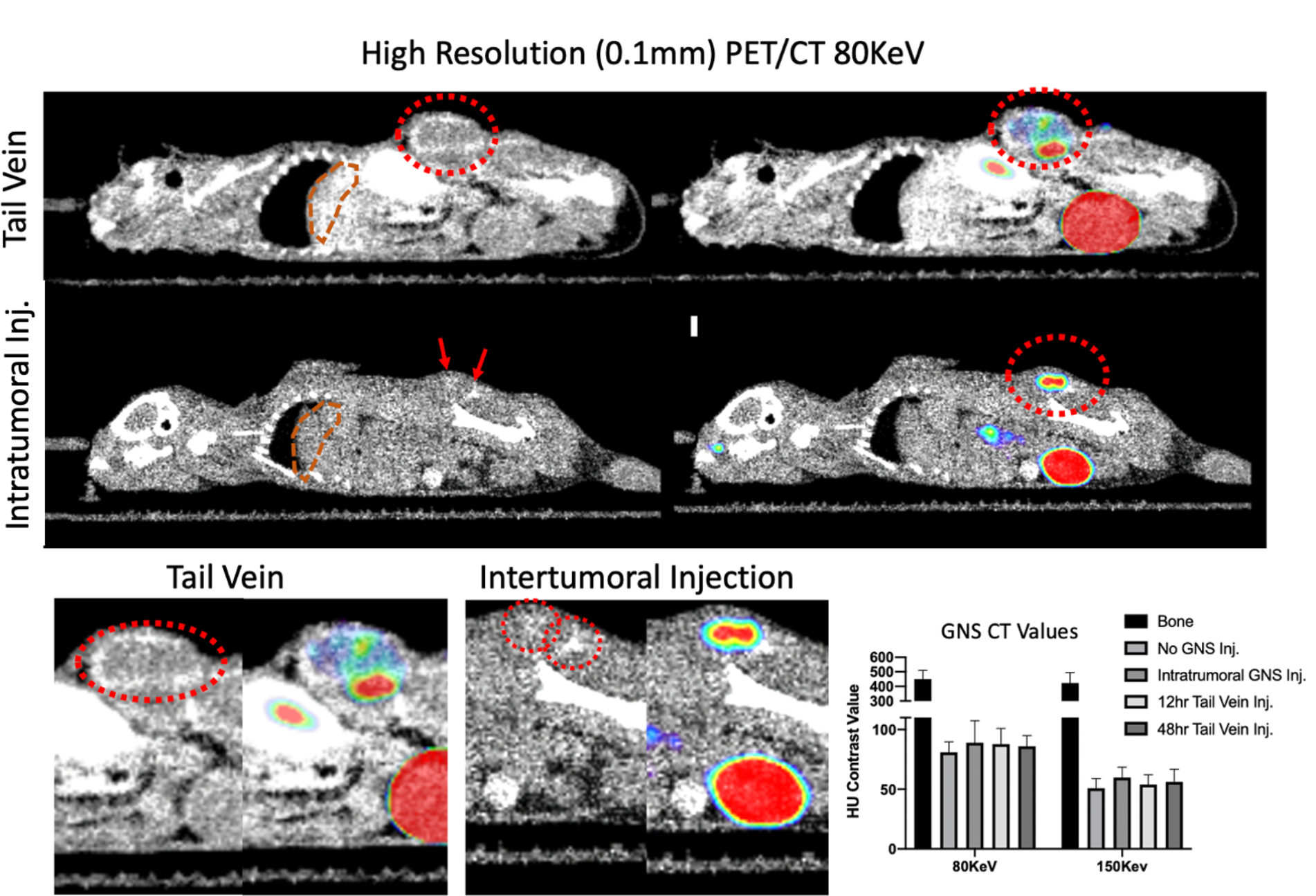
CT imaging of ICG-GNS localization: Compares tail vein injection vs intratumoral injection of ICG-GNS at 80KeV and 150KeV beam energies. Tail vein injected tumors demonstrate increased contrast around the tumor edge, while intratumorally injected tumors show high contrast at ICG-GNS injection sites within the tumor. Although tumors ICG-GNS are elevated in contrast, the results are not significant.

### Radiation therapy and tumor growth dynamics

Compared to SCC7 mice that received no ICG-GNS injection, mice that received either intertumoral injection or tail-vein had a 60% reduction in tumor volume at the end of 30 days post-surgical debulking and fractionated therapy (p<0.05). Mice that received fractionated therapy alone had 50% larger tumors then mice that received intertumoral or tail-vein ICG-GNS administration (p <0 .05). No statistically significant difference in tumor volume was observed between the 4x2Gy and 4x4Gy ICG-GNS injected groups at the conclusion of our experiment (**Figure 5**).

**Figure 5:**
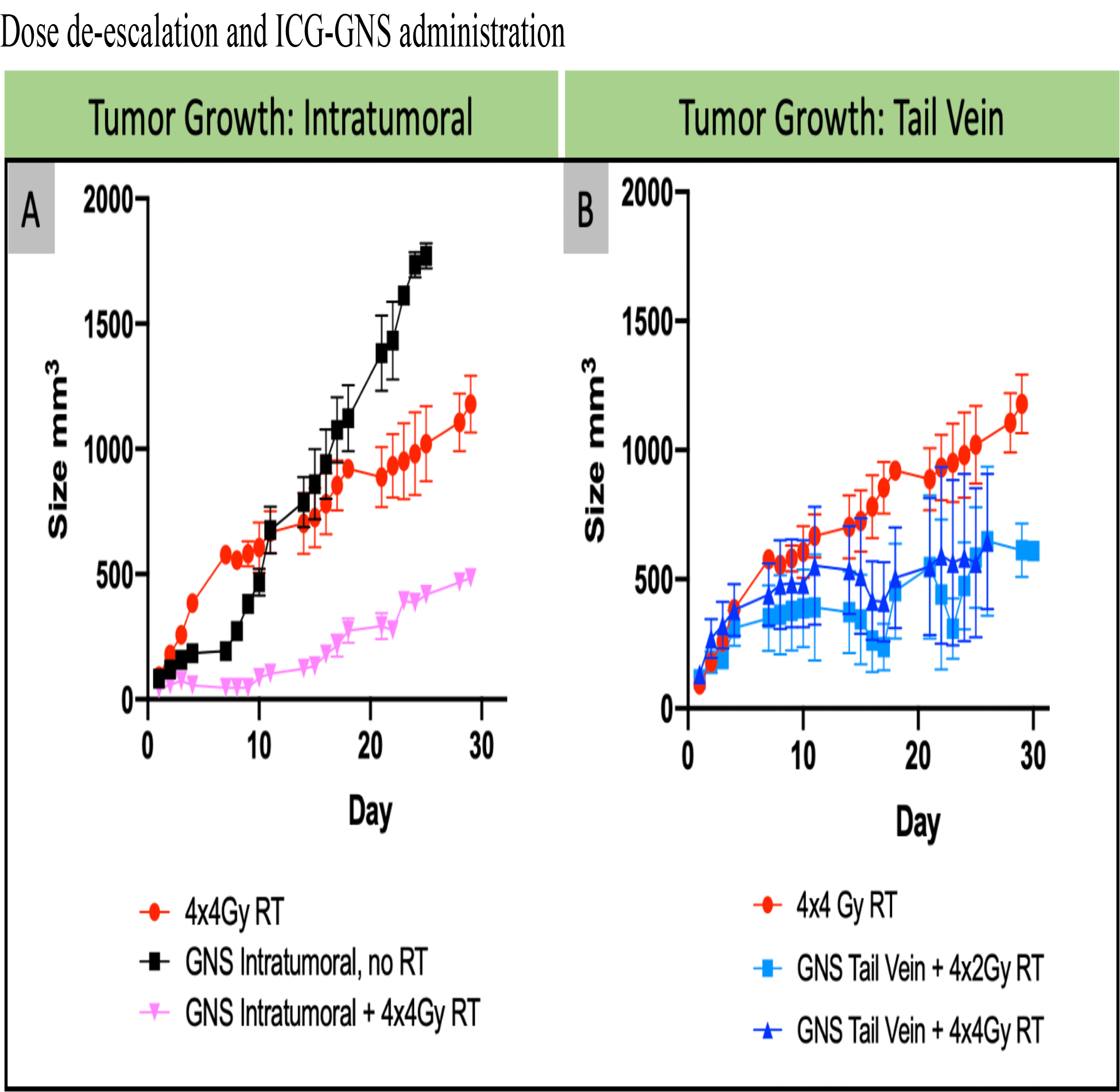
Demonstrates the effectiveness of dose de-escalation methods on intratumorally and tail vein injected ICG-GNS. The black line shows that ICG-GNS alone without radiotherapy had the highest tumor growth rate, while intratumoral injection with 4x4Gy fractionated therapy has the slowest growth rate. At the conclusion of 30 days post-surgery/RT, tail-vein and intramoral injection showed similar tumor sizes around 500cm^2^.

## DISCUSSION

Nanomedicine has attracted increasing attention in recent years because it offers great promise to provide precision diagnostics and personalized medical and surgical treatment. Similarly, real-time cancer margin detection is of great interest for surgical based cancer treatments. Our group has previously demonstrated that DOCI imaging with a 365nm light source provides valuable cancer margin delineation in vivo and intraoperatively.^5-9^ All eukaryotic and prokaryotic cells exhibit an intrinsic natural autofluorescence (AF) easily excited at 365nm, due to the presence of various fluorescent cellular structural components and metabolites, such as flavins, nicotinamide-adenine dinucleotide (NAD), aromatic amino acids, lipofuscins, advanced glycation end products, and collagen.^23,24^ This difference in AF cellular bioproducts between tissues gives distinctive lifetime values that provide natural contrast between cancer and non-cancer tissue when imaged. However, a limitation of 365nm light source is its superficial depth of penetration within tissue as depicted in **Figure 3**. Given that it would be advantageous for surgeons to possess precise tumor depth information when performing a complete oncologic resection, our team sought to utilize ICG-GNS to enhance DOCI capability.

Previously, we have shown that GNSs may be used as a contrast agent and can be functionalized with a wide variety of molecules suitable for surface-enhanced Raman spectroscopy (SERS), positron emission tomography (PET) imaging, and two-photon photoluminescence imaging.^11-13,17,19^ In this proof-of-concept study, not only have we demonstrated that the use of GNSs is suitable with in-vivo DOCI imaging in the near-infrared region, but we have also established that GNS allows for remarkable tumor margin delineation in addition to the capability of assessing tumor depth (**Figure 2,3**). This ICG-GNS nanoparticle platform expands the current DOCI imaging capabilities. The fluorescence intensity spectra for 365nm-excited tissues does not overlap with the 785nm excitation of gold nanoparticles.^25,26^ With the advent of multi-wavelength excitation, DOCI not only distinguishes different tissue types at 365nm, but also adds tumor depth information by way of ICG-GNS emission at 785nm excitation. Such tools will improve cancer resection and aid surgeon in achieving oncologically complete resections. The utility of GNS is shown via both tail vein injection and direct tumoral injections as shown in Figure 4, indicating that clinicians would have the ability of performing either form of GNS administration with adequate accumulation within tumors. As DOCI has already reliably differentiated head and neck SCC from surrounding normal tissue in prior experiments,^5-7,9^ addition of GNSs in delineating deep margins as highlighted in Figures 2 and 3 will allow for improved cancer resection in real time, guiding surgeons to determine suspicious microscopic regions more accurately. Our murine model results indicate that implementation of GNS has the capability to revolutionize DOCI’s clinical promise in cancer care.

Histological evaluation of tumor specimens, detailed in **Figure 2**, identifies the preferential ICG-GNS uptake within the tumor beds at both 4mm and 60μm magnifications. This is seen in both tail vein injected mice and intratumorally injected mice, and despite the assumption that some ICG-GNS is cleared with formalin fixation and tissue processing for H&E slide generation. Tail vein injected ICG-GNS showed ICG-GNS accumulation around nucleated cells at the periphery of the tumor, and tumor vascular sites, while intratumorally injected tumors accumulated larger clumps of ICG-GNS throughout the tumor mass. Furthermore, surrounding muscle in the tumor bed after removal of tumor did not appear to have visible ICG-GNS upon histological evaluation. The histologic findings in this study stand to confirm the DOCI imaging results. The histological slides indicate that GNS selectively accumulate within tumors but not surrounding healthy tissues, likely due to the enhanced permeability and retention (EPR) effect. ^11, 17, 18^

The localization of ICG-GNS in DOCI images, histological slides and CT imaging, detailed in **Figure 2,4**, appears to be consistent. In **Figure 2**, tail vein ICG-GNS injected mice showed a faint boundary around the tumor, while intratumoral administration had a more uniform DOCI value throughout the image. This phenomenon is also seen in CT imaging in **Figure 4**, where tail vein injected tumors have increased contrast on the tumor edge, while intratumorally injected tumors show high contrast at ICG-GNS injection sites within the tumor. This suggests that either modality of ICG-GNS administration and use should be dependent on the clinical context. DOCI, unlike single beam CT, is not concentration dependent, as the fluorescence intensity of a compound does not change its inherent lifetime value. As such, DOCI is limited by its ability to effectively excite a fluorophore.

Materials with high atomic number, such as gold, benefit from the additional ionization at soft X-ray (keV energies) due to the overproduction of photoelectric electrons at those energies. X-rays delivered at the orbital shell binding energies produce further ionization enhancement with characteristic x-rays delivered locally. However, soft x-ray sources are not typically used for cancer treatment (treatment beams 4-25MeV) as their dose delivery remains superficial, disproportionately applying dose to shallow tissue as opposed to deep tumors. However, in the context of head and neck cancers where tumors are predominantly superficial, a soft X-ray treatment approach may benefit from enhance dose delivery of localized GNS. 80KeV is the releasing energy of the K^th^-shell orbital for gold. Gold, when compared to soft tissue provides approximately 100 times more ionization events at 80KeV over soft tissue.^27^ This explains the reduction in tumor growth displayed in mice that received some form ICG-GNS administration mice in **Figure 5**, a finding that is consistent with our prior studies.^13^ This finding is highly clinically significant as it indicates that administration of GNS may allow for radiation dose de-escalation, which is highly advantageous and leads to improved patient quality of life and patient satisfaction. Interestingly, CT imaging at 80KeV to observe the enhanced effect showed either high contrast hot spot at injection sites for intratumorally injected mice or contrast enhancement on the tumor edge for tail vein injected mice. In either ICG-GNS injection scenarios, tumors showed increase contrast over tumors that received no ICG-GNS. Upon assessment of the ROI volumetrically for the entire tumor the CT HU values, we found that HU values were not significantly increased in ICG-GNS injected mice compared to untreated mice. This may be due to the overall lower contrast uptake within the tumors and the differences in ICG-GNS uptake dependent on administration method. In future studies, we plan to implement use of dual energy CT and photon counting CT in order to provide improved image contrast for low concentration contrast agents.

## CONCLUSION

Our study indicates that integration of an ICG/GNS-enhanced DOCI differentiation of tissues could be key to achieving negative margins intraoperatively in real time, optimizing oncologic outcomes and improving patient overall survival while preserving healthy tissue and decreasing morbidity. Furthermore, the use of ICG-GNS is compatible with excising treatment and therapeutic modalities, with the ability for pre-operative CT imaging enhancement, improved intra-operative DOCI margin delineation, and enhance post-operative RT effectiveness.

## ACKNOWLEDGEMENTS

This research is based upon work supported by the National Institutes of Health (R01DE026654 and R01EB028078-01A1).

## References

1. Mignogna MD, Fedele S, Lo Russo L. The World Cancer Report and the burden of oral cancer. Eur J Cancer Prev. 2004;13(2):139–42. Epub 2004/04/22. doi: 10.1097/00008469-200404000-00008. PubMed PMID: 15100581.

2. Shapiro M, Salama A. Margin Analysis: Squamous Cell Carcinoma of the Oral Cavity. Oral Maxillofac Surg Clin North Am. 2017;29(3):259–67. Epub 2017/07/16. doi: 10.1016/j.coms.2017.03.003. PubMed PMID: 28709529.

3. Binahmed, A., Nason, R. W. & Abdoh, A. A. The clinical significance of the positive surgical margin in oral cancer. Oral Oncol. 10.1016/j.oraloncology.2006.10.001 (2007).

4. Zanoni DK, Migliacci JC, Xu B, Katabi N, Montero PH, Ganly I, Shah JP, Wong RJ, Ghossein RA, Patel SG. A Proposal to Redefine Close Surgical Margins in Squamous Cell Carcinoma of the Oral Tongue. JAMA Otolaryngol Head Neck Surg. 2017;143(6):555–60. Epub 2017/03/10. doi: 10.1001/jamaoto.2016.4238. PubMed PMID: 28278337; PMCID: PMC5473778.

5. Tajudeen BA, Taylor ZD, Garritano J, Cheng H, Pearigen A, Sherman AJ, Palma-Diaz F, Mishra P, Bhargava S, Pesce J, Kim I, Sebastian C, Razfar A, Papour A, Stafsudd O, Grundfest W, St John M. Dynamic optical contrast imaging as a novel modality for rapidly distinguishing head and neck squamous cell carcinoma from surrounding normal tissue. Cancer. 2017 Mar 1;123(5):879–886. doi: 10.1002/cncr.30338. Epub 2016 Oct 20. PMID: 27763689.

6. Kim IA, Taylor ZD, Cheng H, Sebastian C, Maccabi A, Garritano J, Tajudeen B, Razfar A, Palma Diaz F, Yeh M, Stafsudd O, Grundfest W, St John M. Dynamic Optical Contrast Imaging. Otolaryngol Head Neck Surg. 2017 Mar;156(3):480–483. doi: 10.1177/0194599816686294. Epub 2017 Jan 24. PMID: 28116982.

7. Pellionisz PA, Badran KW, Grundfest WS, St John MA. Detection of surgical margins in oral cavity cancer: the role of dynamic optical contrast imaging. Curr Opin Otolaryngol Head Neck Surg. 2018 Apr;26(2):102–107. doi: 10.1097/MOO.0000000000000444. PMID: 29517537; PMCID: PMC5846197.

8. Hu Y, Han AY, Huang S, Pellionisz P, Alhiyari Y, Krane JF, Shori R, Stafsudd O, St John MA. A Tool to Locate Parathyroid Glands Using Dynamic Optical Contrast Imaging. Laryngoscope. 2021 Oct;131(10):2391–2397. doi: 10.1002/lary.29633. Epub 2021 May 27. PMID: 34043240.

9. Tam, K., Alhiyari, Y., Huang, S. et al. Label-free, real-time detection of perineural invasion and cancer margins in a murine model of head and neck cancer surgery. Sci Rep 12, 12871 (2022). 10.1038/s41598-022-16975-w

10. Rühle A, Grosu AL, Nicolay NH. De-Escalation Strategies of (Chemo)Radiation for Head-and-Neck Squamous Cell Cancers—HPV and Beyond. Cancers (Basel). 2021;13(9):2204. doi:10.3390/cancers13092204

11. Liu Y, Ashton JR, Moding EJ, Yuan H, Register JK, Fales AM, Choi J, Whitley MJ, Zhao X, Qi Y, Ma Y, Vaidyanathan G, Zalutsky MR, Kirsch DG, Badea CT, Vo-Dinh T. A Plasmonic Gold Nanostar Theranostic Probe for In Vivo Tumor Imaging and Photothermal Therapy. Theranostics. 2015 May 23;5(9):946–60. doi: 10.7150/thno.11974. PMID: 26155311; PMCID: PMC4493533.

12. Weissleder R. A clearer vision for in vivo imaging. Nat Biotechnol. 2001; 19:316–7.

13. Yuan H, Khoury CG, Hwang H, Wilson CM, Grant GA, Vo-Dinh T. Gold nanostarssurfactant-free synthesis, 3D modelling, and two-photon photoluminescence imaging. Nanotechnology. 2012, 23, 075102.

14. Yuan H, Wilson CM, Xia J, Doyle SL, Li S, Fales AM, Liu Y, Ozaki E, Mulfaul K, Hanna G, Palmer GM, Wang LV, Grant GA, Vo-Dinh T. Plasmonics-enhanced and optically modulated delivery of gold nanostars into brain tumor. Nanoscale. 2014 Apr 21;6(8):4078–82. doi: 10.1039/c3nr06770j. Epub 2014 Mar 11. PMID: 24619405; PMCID: PMC4343032.

15. Liu Y, Yuan H, Kersey F, Register J, Parrott M, Vo-Dinh T. Plasmonic gold nanostars for multi-modality sensing and diagnostics. Sensors. 2015; 15:3706–20.

16. Hu, Y. et al. Design and validation of an intraoperative autofluorescence lifetime imaging device. Imaging Therap. Adv. Technol. Head Neck Surg. Otolaryngol. 11213, 23. 10.1117/12.2560000 (2020).

17. Liu Y, et al. Non-invasive sensitive brain tumor detection using dual-modality bioimaging nanoprobe. Nanotechnology; 30:275101.

18. Liu Y, Synergistic Immuno Photothermal Nanotherapy (SYMPHONY) for the treatment of unresectable and metastatic cancers. Scientific Reports. 2017; 7:8606.

19. Chorniak E, et al. Intravital optical imaging for immune cell tracking after photoimmunotherapy with plasmonic gold nanostars. Nanotechnology. 2022; 33: 475101.

20. Singh M, Lima A, Molina R, Hamilton P, Clermont AC, Devasthali V. et al. Assessing therapeutic responses in Kras mutant cancers using genetically engineered mouse models. Nat Biotechnol. 2010;28:585–93

21. Prabakaran PJ, Javaid AM, Swick AD, et al. Radiosensitization of adenoid cystic carcinoma with MDM2 inhibition. Clin Cancer Res. 2017;23(20):6044–6053. doi:10.1158/1078-0432.CCR-17-0969

22. Cosper PF, Abel L, Lee YS, et al. Patient Derived Models to Study Head and Neck Cancer Radiation Response. Cancers (Basel). 2020;12(2):419. doi:10.3390/cancers12020419

23. Surre J, Saint-Ruf C, Collin V, Orenga S, Ramjeet M, Matic I. Strong increase in the autofluorescence of cells signals struggle for survival. Sci Rep. 2018 Aug 14;8(1):12088. doi: 10.1038/s41598-018-30623-2. PMID: 30108248; PMCID: PMC6092379.

24. Croce, A. C. & Bottiroli, G. Autofluorescence spectroscopy and imaging: a tool for biomedical research and diagnosis. Eur J Histochem 58, 2461 (2014).

25. Aldosari FMM. Characterization of Labeled Gold Nanoparticles for Surface-Enhanced Raman Scattering. Molecules. 2022;27(3):892. doi:10.3390/molecules27030892

26. Anzalone A, Gabriel M, Estrada LC, Gratton E. Spectral Properties of Single Gold Nanoparticles in Close Proximity to Biological Fluorophores Excited by 2-Photon Excitation. PLoS One. 2015;10(4):e0124975. doi:10.1371/journal.pone.0124975

27. López-Valverde JA, Jiménez-Ortega E, Leal A. Clinical Feasibility Study of Gold Nanoparticles as Theragnostic Agents for Precision Radiotherapy. Biomedicines. 2022;10(5):1214. doi:10.3390/biomedicines10051214

